# A Normative Model Reveals Functional Optimization of Synaptic Nano-Architecture

**DOI:** 10.64898/2026.09.25.754349

**Authors:** Camille Gontier, Katharina Schmidt, Martín Baccino-Calace, Wayan Gauthey, Markus Sauer, Sina Wäldchen, Divya Sachidanandan, Robert J. Kittel, Martin Müller, Jean-Pascal Pfister

## Abstract

Super-resolution microscopy approaches have revealed that synaptic proteins are not randomly distributed within individual synapses. Rather, they form sub-synaptic clusters that may be coordinated between the synaptic compartments, thereby forming transsynaptic nanocolumns. However, the functional significance of this nano-architecture remains under debate. In particular, it is unclear whether the positions of synaptic proteins optimize a specific cost function. Here, we present a normative modeling framework that integrates synaptic nano-architecture imaged with STED and dSTORM microscopy with synaptic transmission recorded with electrophysiology at the *Drosophila* neuromuscular junction (NMJ). Model-based quantification showed that clusters of Unc13A, which likely demark presynaptic release sites, and postsynaptic glutamate receptors (GluRs) adopt aligned ring-like arrangements in individual synapses. We developed an analytical description of GluR activation relative to presynaptic release position, enabling to link super-resolution imaging data with recorded postsynaptic current amplitudes without relying on costly Markov chain Monte Carlo (MCMC) simulations. Our model predicts that glutamate released from a single vesicle spreads across the entire receptor population. Given this spread, the postsynaptic current amplitude is maximized when glutamate is released close to the active zone (AZ) center, in contradiction to the observed offset of Unc13A clusters. However, the observed relative positions of Unc13A and GluR clusters maximize the postsynaptic current amplitude and minimize the delay between vesicle release and receptor activation during multivesicular release. Hence, transsynaptically-aligned Unc13A-GluR rings optimize for multivesicular but not univesicular release. Given that multi-vesicular release is mainly observed during enhanced presynaptic release at the *Drosophila* NMJ, transsynaptic nano-architecture may be adapted to elevated presynaptic release states.

**Significance statement:** There is evidence that clusters of synaptic proteins, including neurotransmitter receptors, align across the synaptic cleft. However, the functional role of this organization is debated, and it is unclear whether the positions of synaptic proteins allow the optimization of specific cost functions. Here, we developed a normative approach to relate synaptic nano-architecture with synaptic physiology at the *Drosophila* neuromuscular junction. Our approach suggests that the observed positions of presynaptic Unc13A clusters, which may represent presynaptic release sites, and postsynaptic neurotransmitter receptors maximize the amplitude of postsynaptic currents. Hence, synaptic nano-organization likely provides the structural basis for synaptic strength during multivesicular release.

## 1 Introduction

Information transfer at chemical synapses follows a stereotypical process across species: an action potential (AP) triggers presynaptic calcium influx, followed by the fusion of synaptic vesicles with the presynaptic membrane. The liberated neurotransmitter molecules then diffuse into the synaptic cleft and bind to postsynaptic neurotransmitter receptors, thereby increasing their open probability, and, in case of ionotropic receptors, producing an ionic current. The receptor activation probability depends on neurotransmitter concentration [1, 2], and the neurotransmitter concentration is not uniform but follows a time-dependent gradient from the release site [3, 4]: synaptic functions thus depend on the relative distances between release sites and receptors [5–7], and more generally on the geometry of synaptic proteins [8].

Superresolution microscopy allows to observe both presynaptic Unc13A (proteins involved in the spatial organization of the synapse, and used as a proxy for the position of neurotransmitter release sites [9]) clusters, and postsynaptic glutamate receptors (GluR) clusters. Interestingly, the positions of these proteins across the synaptic plane have been observed not to be random, but to follow specific patterns: there is indeed evidence for synaptic [10–12] and transsynaptic nano-architecture [13]. More specifically, super-resolution imaging of presynaptic release suggests that vesicles fuse close to presynaptic RIM clusters: based on the observation that presynaptic RIM clusters align with postsynaptic PSD95 clusters, these data support a model of synaptic transmission within transsynaptic nano-columns [13]. Furthermore, data recorded at the *Drosophila* neuromuscular junction (NMJ) also suggest a higher-order, ring-like organization of transsynaptic nano-architecture [14].

However, the relationship between this transsynaptically-aligned nano-architecture and synaptic physiology remains elusive. Presynaptic nano-architecture shapes presynaptic release under baseline conditions [10–12] and synaptic plasticity [15]. Given that receptor open probability decreases with distance to the presynaptic release site [1–4], it has been hypothesized that transsynaptic nano-architecture optimizes synaptic transmission by aligning presynaptic release sites and postsynaptic receptors. In line with this hypothesis, disruption of transsynaptic nano-alignment impairs synaptic transmission [8, 16]: a previous study [16] observed effects on the amplitude of synaptic miniature transmission, while another [8] did not, and instead revealed altered AP-evoked synaptic transmission.

Moreover, there has been no systematic investigation of whether higher-order organization of transsynaptic nano-alignment maximizes specific cost functions. Although models of synaptic transmission have been widely used to study how synaptic structures impact information transmission, and especially how they affect the peak postsynaptic current [5, 17, 18], they have not been confronted to the actual organization of biological synapses. In particular, the position of presynaptic Unc13A clusters and of postsynaptic receptors that maximizes synaptic currents amplitudes (or minimizes their onset delay) in simulations has not been compared to their actual positions. In line with previous reports of optimality in neural systems [19, 20], our main contribution is thus to propose a normative approach to the organization of synaptic proteins, and suggest that it might optimize a certain biological cost function (e.g. the mean current amplitude or its time delay after vesicle release).

The main novelty of our approach is to combine microscopy observations of the position of presynaptic release sites and of postsynaptic receptors, and electrophysiology recordings of spontaneous postsynaptic currents: this allows to confront synaptic transmission (i.e. the distribution of postsynaptic events) to its geometry (i.e. the spatial distribution of proteins). We applied this approach to the *Drosophila* NMJ, a structure that is well established for studying molecular mechanisms of synaptic transmission and has been less studied than synapses in the CNS.

Our first contribution was to employ a theoretical approach to confirm whether preand postsynaptic nano-clusters indeed arrange in ring structures and are transynaptically organized (Section 2.1). The positions of Unc13A and GluRs clusters on the synapse can be modeled as a Gibbs Point Process (GPP), which is characterized by two elements: its intensity function (i.e. the spatial probability density of points), and its interaction function (i.e. the spatial dependencies between points’ positions and types). For a given set of observation, the likelihood of different GPPs can be computed, and their goodness-of-fit can be compared. The tested model yielding the highest goodness-of-fit is the one with ring-like intensity functions (i.e. Unc13A and GluRs being organized in ring structures and not randomly scattered across the synapse); and with type-specific interaction functions (i.e. Unc13A and GluRs being likely to be aligned), which is in line with historical observations [13, 14].

Our second contribution was to derive a phenomenological model for the sensitivity of postsynaptic glutamate receptors, i.e. the probability that a receptor will be activated by glutamate released at a distance *r* (Section 2.2). The main shortcoming of previous *in silico* computational models of synaptic transmission was their reliance on lengthy simulations in which the position of each molecule as well as the current state of each receptor is tracked, which makes them computationally challenging. Starting from the heat kernel (an analytical solution to the heat equation) to model the neurotransmitter concentration as a function of time and distance to the release site, we derived a mean-field model of the sensitivity of postsynaptic receptors: this approach does not require to simulate a Brownian motion, nor to specify many free parameters [8].

The obtained activation probability is a function of the distance *r* between the release site and the receptor, but also of the neurotransmitter diffusion coefficient *r*_0_, which is a free parameter of the model. A small value for *r*_0_ indicates that only receptors in the vicinity of the release site will be activated, while a large value indicates the neurotransmitters from a single release site are likely to cover a large area of the postsynaptic density (PSD). Hence, how the geometry of synaptic proteins impacts synaptic transmission critically depends on the value of *r*_0_. Our third contribution was thus to obtain a point estimate of *r*_0_ by computing its likelihood given a set of recorded synaptic currents (Section 2.3).

Our final contribution was to compare the observed positions of synaptic proteins to their theoretically optimal ones. In the univesicular case (Section 2.4), given the estimated value for the neurotransmitter diffusion coefficient *r*_0_, the glutamate release site position that maximizes the postsynaptic current amplitude and minimizes its time-to-peak is at the center of the AZ. However, the density of observed Unc13A positions peaks approximately 100nm away: this offset between the theoretically optimal and empirically observed positions of release sites can be partially resolved by considering structural constraints and the presence of other proteins which may contribute to excluding synaptic vesicles from the center. On the other hand, release from observed Unc13A locations is predicted to optimize for postsynaptic amplitude during multivesicular release (Section 2.5): we computed the theoretically optimal position of presynaptic release sites and showed it closely matches their actual observed positions. We finally closed the loop, and show that these theoretically optimal positions yield the same results as actual observed positions when performing the model selection task from our first contribution.

## 2 Results

### 2.1 Unc13A and glutamate receptor clusters organize in transsynaptic nanocolumn rings

The *Drosophila* NMJ has emerged as one of the best model systems to investigate synaptic nano- architecture and synaptic transmission [21]. Presynaptic protein clusters are arranged around a voltage-gated calcium channel cluster at the AZ center [10–12]. Moreover, presynaptic protein cluster rings align with postsynaptic receptor clusters, and were suggested to form transsynaptically-aligned nanocolumn rings [14]. However, synaptic nano-architecture has not been analyzed quantitatively with models assuming specific protein distributions. Our first step was therefore to derive a quanti- tative description of synaptic protein distributions using a model-based approach. To this end, we detected presynaptic Unc13A clusters, which are thought to represent presynaptic release sites [9, 15], and postsynaptic GluR clusters (anti-GluRIIC) in maximum intensity projection STED data (Fig. 1A). The number and dimensions of the protein clusters are limited by the STED resolution, and we detected the peak of each cluster using local fluorescence intensity gradient analysis [14]. A total of 365 active zones from 5 NMJs were analyzed. The average number of Unc13A clusters per AZ was 2.61 *±* 1.21, while the average number of GluR clusters per PSD was 4.06 *±* 1.17. The average distance of protein clusters from the AZ center was 108 *±* 28*nm* for Unc13A and 121 *±* 21*nm* for GluR (Table 1).

**Table 1:**
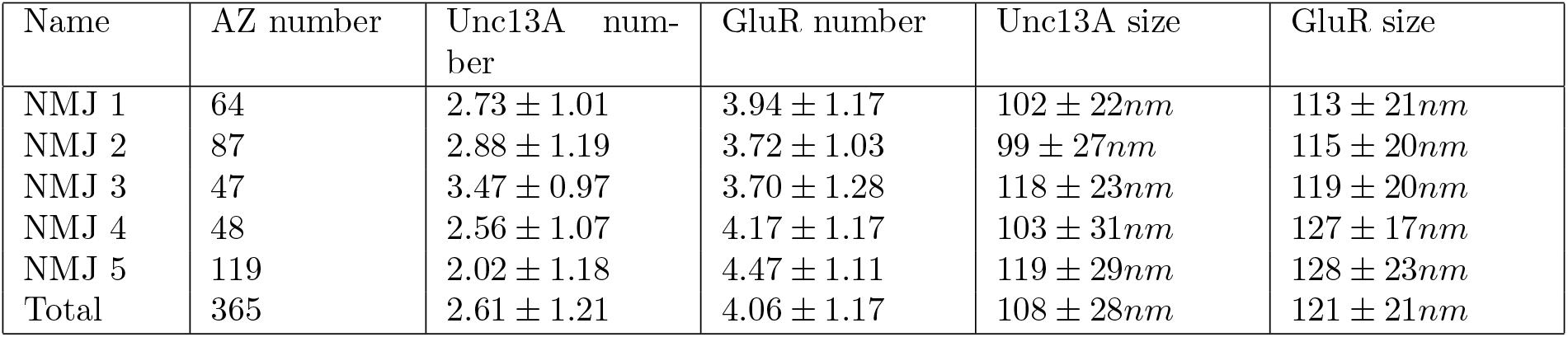
Geometries of active zones for different observed NMJs.

**Figure 1.**
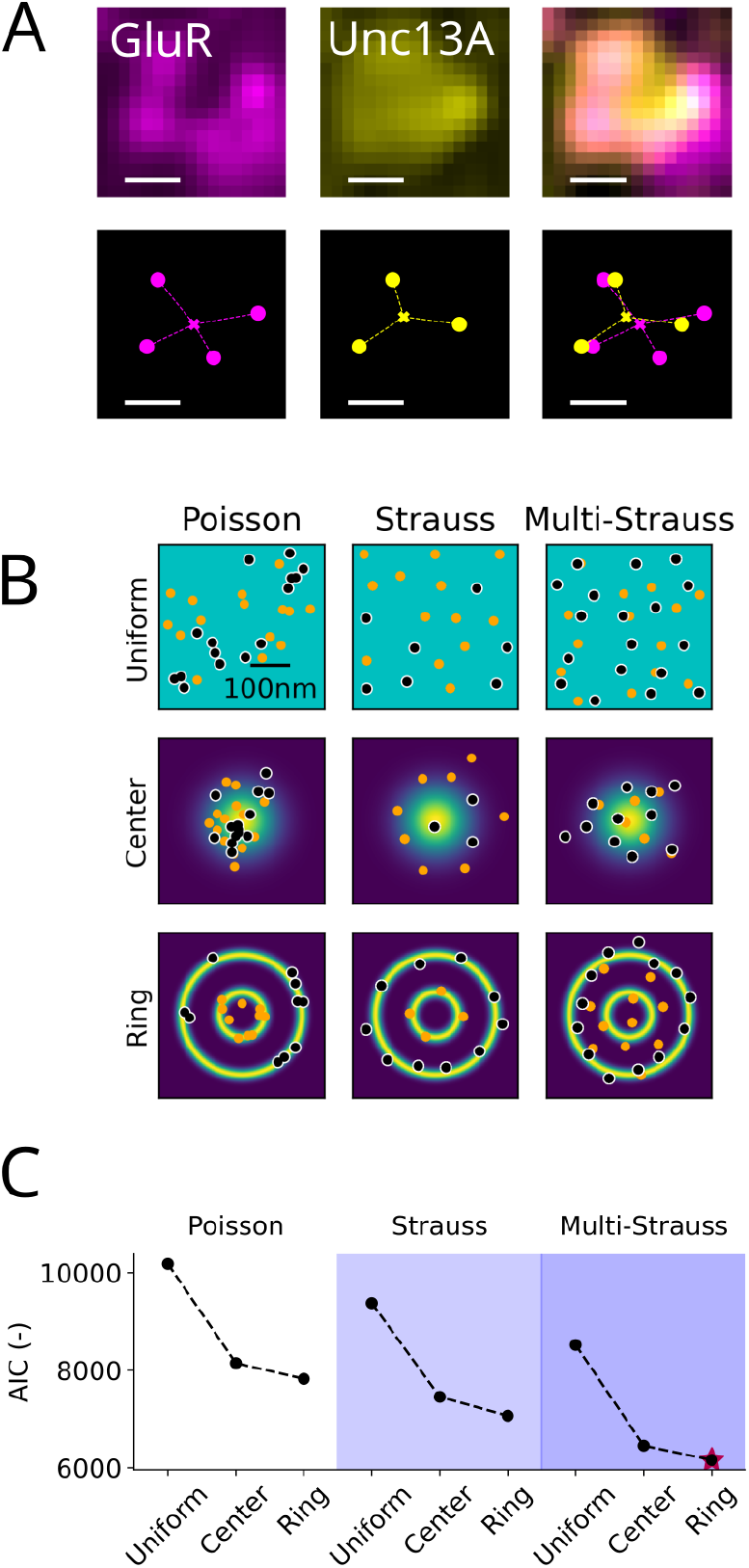
Unc13A and glutamate receptor clusters organize in transsynaptic nanocolumn rings. A: Maximum intensity projection STED image (top) and detected clusters (circles) and centers (crosses) (bottom) of anti-GluRIIC (‘GluR’, left) and anti-Unc13A (middle) of a representative *Drosophila* NMJ synapse. Scale bar: 100nm. B: A model is defined by its intensity function (*Uniform, Distance to center*, or *Ring patterns*), and by its pairwise interaction function (*Poisson, Strauss*, or *Multi-Strauss*). These three intensity functions and three interaction functions define nine competing models which are compared based on their ability to fit the observations. Orange and blue circles demark GluRIIC and Unc13A cluster locations, and localization probability is color coded from blue (low) to yellow (high). C: Results of model fitting for a representative NMJ. A lower AIC indicates a better fit. The model with the lowest AIC has an intensity function structured in ring-like patterns, and the best model of interaction is a multi-Strauss process. Results for other NMJs are displayed in Supp. Fig. 1.

The position of Unc13A and GluR clusters can be understood as a multitype point process, where each point is assigned a position (*x*_1_, *x*_2_) on the 2-dimensional AZ plane as well as a type (either *Unc13A* or *GluR*). We proposed several models for this point process, each characterized by distinct intensity functions and pairwise interactions, and conducted model selection to evaluate their goodness-of-fit (see Gibbs Point Processes). Finding out which model fits the data best allows to draw conclusions on cluster distribution and interaction. First, point processes are characterized by their intensity function (i.e. the density of points), which can be either uniform or a function of the position in the 2D plane for each type of points. We proposed three different intensity functions: a uniform one (i.e. the point density is the same over the whole synapse surface, top row in Fig. 1B); a distance-to-center intensity function (the point density decreases as a function of the distance to the synapse center, middle row in Fig. 1B); and a ring pattern intensity function (lower row in Fig. 1B).

Second, point processes are characterized by the interpoint interactions, i.e., by possible dependencies between the points’ positions and types. Here, we proposed three different interaction schemes: a Poisson point process, in which all points are independent, irrespective of their types (left column in Fig. 1B); a Strauss point process, which specifies an exclusion radius between two points (middle column in Fig. 1B); and a multi-Strauss point process, which further generalizes the Strauss process to be type-dependent. More specifically, in a multi-Strauss point process, the interaction radius and the parameter *γ* controlling the probability of finding a point in the vicinity of another will depend on the respective types of points (*Unc13A-to-Unc13A, Unc13A-to-GluR*, or *GluR-to-GluR*).

These three intensity functions and three pairwise interaction functions thus define nine possible generative models (Fig. 1B), which were compared based on their respective Akaike Information Criterion (AIC) for the microscopy data (see Model comparison). The AIC is a model selection score that compares candidate models based on their ability to explain observations while penalizing their number of free parameters [22–24]: the lower the AIC, the better the model.

The AIC scores for the different models are compared in Fig. 1C. The model with the best AIC has an intensity function structured in ring-like patterns, in line with previous observations of ring-like patterns of Unc13A [15] and GluRs [14]. Furthermore, the best model of interaction is a multi-Strauss process: protein clusters are neither independent, nor have a simple exclusion radius. Rather, Unc13A and GluR clusters have a tendency to be aligned: the inferred interaction parameters *γ* for the multi-Strauss model are given by *γ*_*GluR,GluR*_ = 0.014, *γ*_*Unc*13*A,Unc*13*A*_ = 0.0072, and *γ*_*GluR,Unc*13*A*_ = 1.17, i.e. the within-type interaction parameters (*γ*_*GluR,GluR*_ and *γ*_*Unc*13*A,Unc*13*A*_) are close to zero, indicative of an exclusion radius for protein clusters in the same synaptic compartment; while the between-type interaction parameter *γ*_*GluR,Unc*13*A*_ is larger than 1. This is also in line with previous observations of transsynaptically aligned nanocolumns [13, 14]. Hence, our model comparison suggests that Unc13A and GluR clusters are arranged in transsynaptically-aligned rings.

### 2.2 A fast analytical model predicts GluR activation relative to presynaptic release sites distance

We next aimed at developing an analytical model of GluR activation as a function of the distance between Unc13A and GluR clusters. Previous models relied on Monte Carlo simulations considering the position of each neurotransmitter molecule in the synaptic cleft, as well as the state of each postsynaptic receptor [3, 4, 18, 25]. While these models fit recorded mEPSCs [26], they rely on numerical simulations and are not amenable to mathematical analysis. For instance, the activation probability of a receptor as a function of its distance to the glutamate source can only be computed by averaging several numerical simulations. Moreover, such models have a large number of free parameters (including the rate constants for postsynaptic receptors), which may be hard to accurately estimate from data [3], thereby preventing the model from correctly generalizing to future observations, and leading to identifiably problems. We first derived an analytical formulation for the maximal neurotransmitter concentration at any point of the AZ plane following a release event; then, we proposed a non-linear function to transform this concentration into a postsynaptic receptor activation probability.

The process of mEPSC generation following a spontaneous presynaptic release event involves three main steps [4, 27]. First, the release of glutamate into the synaptic cleft. This release process can be approximated as instantaneous [25], or can be modeled as an exponentially decaying source of glutamate [4]. Second, the diffusion of glutamate within the synaptic cleft can be modeled as a random walk of glutamate molecules. Given the small cleft width (approx. 15.9nm [28]) compared to the AZ surface, it is often approximated as a two-dimensional diffusion process, with an experimentally observed diffusion coefficient *D* [4, 25]. Finally, the glutamate molecules bind to postsynaptic receptors and increase their open probability, thus eliciting a postsynaptic current. Some GluRs are clustered, while others are isolated and scattered across the postsynaptic membrane [5]. GluRs are often modeled as kinetic state diagrams. For instance, in [26], AMPAR kinetics are modeled using a 7-state model which kinetics rate constants are fitted to experimentally recorded mEPSCs. The initial state corresponds to the resting state, and the activated states are reached upon successive binding of several glutamate molecules, followed by a switch to desensitized states, with the transition between receptor states usually following a discrete Markov process [5]. In line with [4], we used the heat equation to represent the neurotransmitter concentration as a function of time and distance to the release site. The heat kernel [4] provides an analytical solution to the 2D heat equation. The glutamate concentration *ρ* at a distance *r* from the release site and at a time *t* after its release is equal to

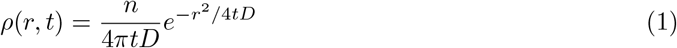

where *n* is the number of glutamate molecules released per vesicle and *D* is a diffusion coefficient expressed in *μm*^2^*/s*. Solving 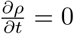 for *t* yields

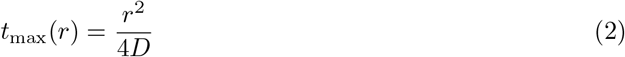

i.e., the time after release at which the maximum glutamate concentration will be reached increases with the square of the distance *r*. Re-injecting Eq. 2 into Eq. 1 finally yields

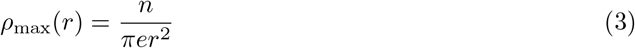

i.e., the maximum glutamate concentration at a distance *r* from the release site will be proportional to 1*/r*^2^. Having derived a link between the distance to glutamate release *r* and the maximal glutamate concentration, our second step was to derive a link between this maximal observed glutamate concentration and glutamate receptor activation probability *p*(*r*). We approximated it as (see Statistical model of glutamate spread and postsynaptic receptor activation for a detailed derivation)

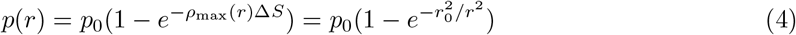

where Δ*S* is the surface of a postsynaptic receptor; *p*_0_ is a scale parameter defining the affinity between glutamate and postsynaptic receptors (a distance *r* = 0 does not necessarily imply that the receptor will be in an activated state [5]); and 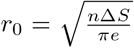 is the effective diffusion radius (i.e. *p*(*r*) *≈ p*_0_ for *r ≤ r*_0_). Note that this model readily extends to multivesicular release (see Extension to multivesicular release). Interestingly, *p* is not a continuously decreasing function of the distance *r*: it plateaus at *p*_0_ for short distances, before decreasing after a cutoff distance close to *r*_0_ (Fig. 3C). This plateau is coherent with the presence of transsynaptic filaments [29] that reduce the diffusion of glutamate in the plane parallel to the active zone, making their diffusion anisotropic [30].

To assess the validity of Eq. 4, we compared it to previous results using numerical simulations to compute the proportion of activated receptors as a function of their distance to a release site. A pharmacodynamical model of AMPA receptors activation, whose kinematic parameters were fitted on recorded mEPSCs, was used in [26]. Using this numerical model, the authors computed the number of open receptors following glutamate release in three different scenarios. The first one simulates one GluR cluster and one release site: the number of activated receptors is computed as a function of the distance between the release site and the cluster (Fig. 2A). The second scenario considers one release site and two GluR clusters, one being located underneath the release site and the second one at a varying distance. The number of activated receptors is computed as a function of this distance (Fig. 2B). Finally, the third scenario simulates one release site flanked by two receptor clusters. The number of activated receptors is computed as a function of the displacement of the release site (Fig. 2C).

**Figure 2.**
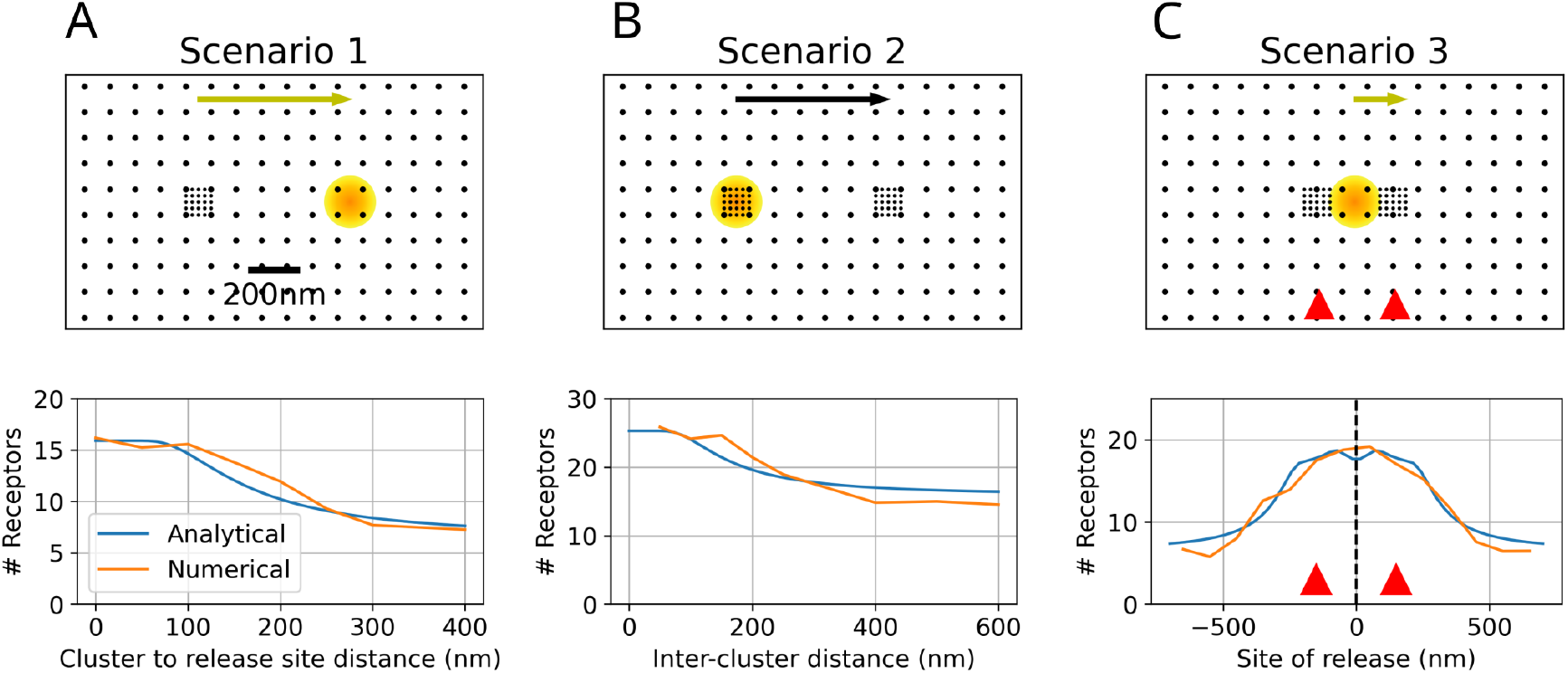
A fast analytical model predicts GluR activation relative to presynaptic release sites distance. The number of activated GluRs (black dots) following glutamate release in one location (orange) is computed in different scenarii. The outputs of our analytical model (Eq. 4, blue) are compared to results from simulations using the complete pharmacodynamical model in [26] (orange). A: 1 release site, 1 GluR cluster, different positions of the release site. B: 1 release site, 2 GluR clusters, different positions of one of the GluR clusters. C: 1 release site, 2 GluR clusters, different positions of the release site.

**Figure 3.**
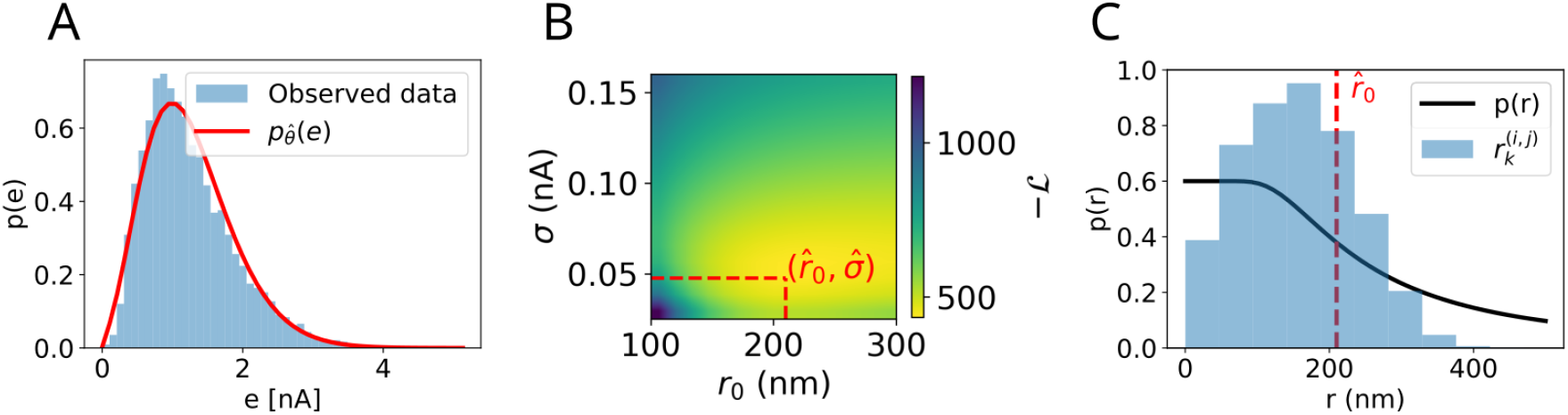
Presynaptic release activates the majority of postsynaptic receptors. A: mEPSC amplitude *e* histogram (blue) and probability distribution 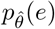 (Eq. 32) after optimal parameter fitting (red). The model provides a good quantitative fit of the observed mEPSC amplitude distribution. B: Negative log-likelihood profile for Eq. 5. The fit between the model 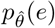 and the observed mEPSCs amplitudes is maximal for large values of *r*_0_ (*>* 150*nm*), indicative of large glutamate spread. C: Value of *p*(*r*) (Eq. 4) for the fitted values of 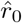 (black), and histogram of the Unc13A-to-GluR distances 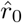 (Eq. 27) observed from all recorded NMJs (blue). Due to the large diffusion radius *r*_0_, most receptors are activated by a single glutamate release event.

The results of simulations based on the numerical pharmacodynamical model in [26] are plotted as orange lines on Fig. 2. Free parameters of our analytical model *r*_0_ and *p*_0_ were fitted by minimizing the difference between the output of Eq. 4 and the results of the numerical simulations: this yielded estimated values for this specific example of *r*_0_ = 142 nm and *p*_0_ = 0.38, which are in line with previous measurements [1, 2, 6, 31]. Results from our simplified analytical model (Eq. 4) after fitting of *r*_0_ and *p*_0_ are plotted in blue, and share a good qualitative similarity with [26]. More generally, the shape of Eq. 4 is in line with results of previous numerical simulations computing receptor activation probability as a function of its distance to a release site (e.g. Fig. 4 in [4]). Our analytical model thus reproduces the experimentally estimated receptor open probability, and circumvents computationally intensive numerical simulations of individual receptor dynamics. Hence, our model allows to systematically explore the relationship between GluR and Unc13A cluster localization and synaptic transmission.

**Figure 4.**
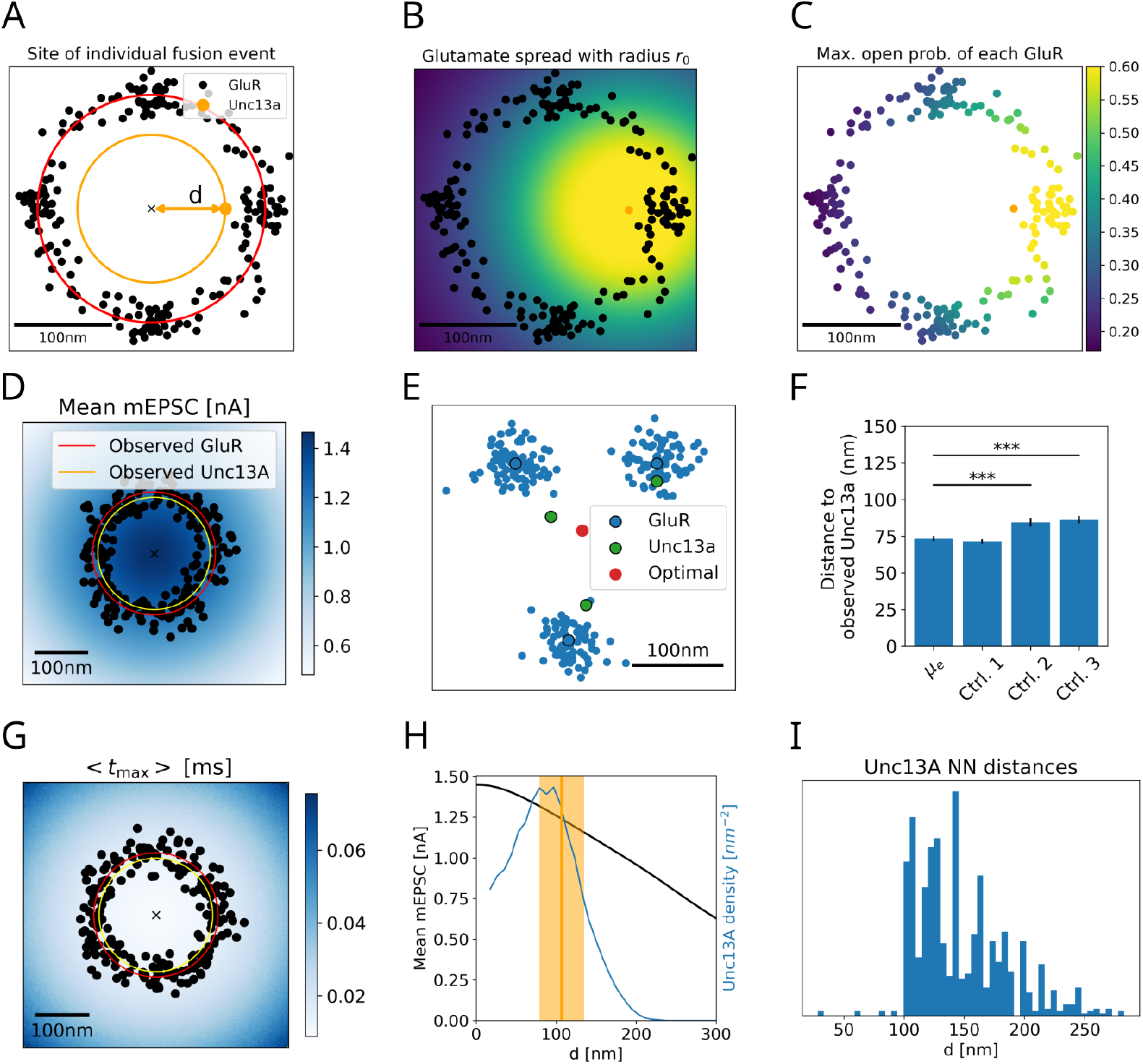
Glutamate release from the AZ center optimizes postsynaptic current amplitude. A-C: Statistical model of the synapse. GluRs (black dots) are randomly located around a ring (red circle) whose radius (121nm) corresponds to the average observed distance between GluRs and the center of AZ. We studied the effect of a simulated fusion event at a distance *d* from the center of the AZ. D: At each point of the AZ, the color code (blue scale) corresponds to the mean postsynaptic current *μ*_*e*_ following glutamate release at this location. *μ*_*e*_ is maximized for a release close to the center. E: Example synapse. The positions of GluRs (blue) was held fixed to their observed values. The optimal position for a release site to optimize *h* (red) was compared to actual Unc13A positions (green). F: The average distance between actual and *μ*_*e*_-optimal release sites was compared to the distance between actual release sites and the center of the AZ (*Ctrl. 1*), a random GluR cluster position (*Ctrl. 2*), and a random position on the AZ disk (*Ctrl. 3*). G: Same as D, but plotting the average activation time *< t*_max_ *>* as a function of the release location. It is minimized for a release close to the center. H: *μ*_*e*_ as a function of the distance *d* between the release site and the AZ center (dark line) vs. observed Unc13A density (blue line). Yellow line: average observed distance to center for Unc13As (shaded area: s.d.). I: Observed density of nearest neighbor distances between Unc13As across all observed AZs.

### 2.3 Presynaptic release activates the majority of postsynaptic receptors

Next, we used our analytical model to relate experimentally observed Unc13A and GluR cluster localizations to postsynaptic current amplitudes in order to estimate glutamate diffusion and the fraction of activated receptors. Whether presynaptic release sites and postsynaptic glutamate receptors are organized to optimize synaptic transmission depends on the spread of glutamate captured by the parameter *r*_0_ in Eq. 4. A low *r*_0_ value is indicative of low glutamate diffusion, and that glutamate receptors should be located close to a release site to have a high activation probability. Conversely, a high *r*_0_ indicates that glutamate diffuses across the entire receptor field, activating receptors even at distances far from the release site.

Although estimates have been previously obtained for other synaptic parameters (e.g. *p*_0_ [1, 2, 6, 31], or the quantal postsynaptic current [1–3, 32]), a reliable estimate for *r*_0_ at the *Drosophila* NMJ is lacking. Our third contribution was thus to obtain a maximum likelihood estimate of *r*_0_ based on a set of observed spontaneous miniature postsynaptic currents (mEPSCs). The likelihood of a given current amplitude is a function of the observed Unc13A and GluR cluster localizations (Section 2.1), of the postsynaptic receptor activation probability (Section 2.2), and the glutamate spread *r*_0_. Fitting our model on the recorded mEPSC amplitude distribution allows to infer the values of the free parameters, and to verify that our model can explain the observed amplitude distribution.

Spontaneous postsynaptic currents had been recorded from *Drosophila* NMJs at a holding potential of *−*100*mV* [33], and values of mEPSCs amplitudes were extracted from raw data using miniML [34]. The distribution of 11482 recorded events is shown in Fig. 3A: their average value is 1.24*nA*, consistent with previously recorded values [35].

We proposed a generative model for mEPSCs (see Generative model of postsynaptic currents): the probability density function for a postsynaptic current amplitude *e* is defined as *p*_*θ*_(*e*), where *θ* = [*μ*_*N*_, *σ*_*N*_, *r*_0_, *p*_0_, *q*_0_, *σ*] is a vector of the parameters of the model: the mean *μ*_*N*_ and standard deviation *σ*_*N*_ of the number of GluR receptors per PSD (Eq. 29), the effective radius *r*_0_ and max. opening probability *p*_0_ (Eq. 4), and the mean *q*_0_ and standard deviation *σ* of the current amplitude elicited by the activation of one GluR (Eq. 30).

Considering biological constraints on the parameters *θ* allows reducing the number of free parameters to be fitted, and hence the risk of overfitting. Similar values for *p*_0_ and *q*_0_ had been obtained in previous studies, i.e. *p*_0_ = 0.6 [1, 2, 6, 31] and *q*_0_ = 12*pA* [1–3, 32]. Furthermore, we estimated the number of GluRs per PSD using direct stochastic optical reconstruction microscopy (dSTORM) [36]. This approach yielded a mean GluR number of *μ*_*N*_ = 214, with a standard deviation *σ*_*N*_ = 64 (see GluR quantification via dSTORM). The distribution was fitted to the observed mEPSCs amplitude distribution 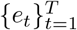 to infer the values of the remaining parameters *r*_0_ and *σ*:

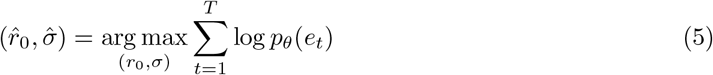

Eq. 5 gave inferred values of 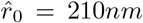 and 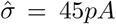 (Fig. 3B), which provide a good qualitative fit to the data (compare the distribution 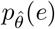to the distribution of observed events in Fig. 3A). Furthermore, the inferred value 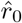 for glutamate spread is large compared to the average PSD diameter (the average GluR diameter being approx. 242 nm, see Table 1), meaning that glutamate is not confined to the vicinity of its release site but reaches most receptors of a given synapse. One can compare the profile of *p*(*r*) (Eq. 4) for 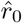 (solid line in Fig. 3C) to the observed distribution of Unc13A-GluR distances (blue histogram in Fig. 3C): approx. 75% of these distances are shorter than 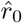. This value is consistent with previous estimates [3, 6, 30].

### 2.4 Glutamate release from the AZ center optimizes postsynaptic current amplitude

Our final contribution was to assess whether the observed organization of release sites and glutamate receptors (Section 2.1), given the sensitivity of postsynaptic receptors (Section 2.2) and the diffusion coefficient of glutamate (Section 2.3) effectively allows to maximize synaptic strength. We first studied the effect of release site locations on univesicular synaptic transmission for a model synapse. It follows from microscopy observations that synapses have on average *μ*_*N*_ = 214 GluR receptors on the postsynaptic density organized in 4 clusters, and that their mean distance to the center of the AZ plane is 121 *±* 21*nm* (Table 1). An average postsynaptic receptor field was thus created by modeling 214 GluRs on a ring structure with a radius of 121nm around the center of the AZ (black dots and red circle in Fig. 4A). We then assessed how different metrics evolve as a function of the position of a single synthetic release site (orange dot in Fig. 4A-C): the GluR activation probability, GluR activation time, and the mean mEPSC amplitude *μ*_*e*_.

Two different regimes will govern the optimal location of release sites depending on the value of *r*_0_: at low *r*_0_ (low glutamate spread), it is optimal for the release sites to be close to the postsynaptic clusters to maximize *μ*_*e*_; at high *r*_0_ (high glutamate spread), it is optimal to be close to the center of the AZ to activate several postsynaptic clusters. Given the large estimated value 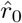 (Fig. 3C), the position of a release site that maximizes the mEPSC amplitude should thus be close to the center of the AZ. As expected, the maximal mean mEPSC is obtained for a release site located at the center of the AZ: *μ*_*e*_ decreases with the distance between the simulated release site and the AZ center (Fig. 4D).

This analysis assumed an ideal model of the synapse, i.e. where the number of proteins and the radius of their ring-structure is equal to the mean observed values. Actually, both the number and position of Unc13A and GluRs are highly variable across synapses (Table 1). To ensure our normative theory is robust to this variability, we compared the observed positions of Unc13A to their theoretically optimal values given the observed position of GluRs. More specifically, for each observed synapse, we fixed the position of GluRs to their observed positions, performed numerical optimization to obtain the position of Unc13A that allows to maximize a given cost function *h*, and ensured these optimal positions are close to their observed ones.

At the *k*-th synapse, the positions of the *M*_*k*_ Unc13A and *N*_*k*_ GluRs are noted 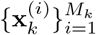 and 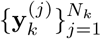 (as in Eq. 27). We also define the function 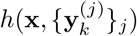 which is the expected postsynaptic current amplitude following glutamate release at a position **x** (Eq. 35). The optimal glutamate release site position to maximize the postsynaptic current amplitude was computed as such:

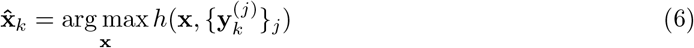

The theoretically optimal position 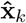 was then compared to its closest neighbor among the observed positions 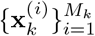. As seen in Fig. 4E for an example synapse, the release site position that optimizes *h* (red) is close to the plane center. This process was repeated across all observed AZs: the distance between the optimal and actual Unc13A positions is not significantly different than the distance between the AZ center and the centermost observed release site (Fig. 4F, *μ*_*e*_ vs. *Ctrl. 1, p* = 0.41). These analyses thus hint at a theoretically optimal position for glutamate release that should be close to the center of the AZ. Moreover, the conclusion still holds even when considering a different cost function, e.g. the average activation time of GluRs *< t*_max_ *>* (Eq. 2), which is also minimized when glutamate is released at the center of the AZ (Fig. 4G).

However, this theoretically optimal position for Unc13A is at odds with their observed distribution, whose mode is not at the center of the AZ but on a ring-like structure approx. 100nm away from it (Fig. 1). The mean postsynaptic current is maximized when the release site is located close to the AZ center, and decreases with the distance between the AZ center and the glutamate release site (Fig. 4H, dark line); but the observed density of Unc13A is maximal at a distance *d ≈* 100nm from the center (Fig. 4H, blue line).

This offset between the normative predicted position and the peak of the observed Unc13A density can be partially explained by structural constraints, and especially by the presence of other proteins at the AZ center which may contribute to excluding synaptic vesicles from the center. The AZ center is indeed occupied by an electron dense structure with the scaffold Bruchpilot (Brp) [12] and by a cluster of voltage-gated calcium channels (VGCC) [10] which is itself surrounded by RIM-binding proteins. Another study [37] had shown freeze-fracture images of putative calcium channels below the electron-dense AZ cytomatrix. This is consistent with previous results showing that the average number of docked synaptic vesicles is low at the AZ center, and that Unc13A levels do not peak at the AZ center, but with an offset of around 70nm [15]. Furthermore, the offset between actual and predicted release site positions could also be linked to an exclusion radius within Unc13A. The distribution of nearest neighbor distances for Unc13As across all AZs shows a sharp increase at approx. 100nm (Fig. 4I), which is coherent with a hard exclusion radius between release sites and with the observed density of Unc13A positions.

Overall, the univesicular theory predicts that the optimal position for a glutamate release site is at the center of the AZ, while the actual Unc13A density peak is observed 100nm away from the AZ center: this offset can be partially explained by structural constraints on presynaptic proteins. Evolution selected for VGCC and other proteins, such as Brp and RIM-binding protein or components of the AZ cytomatrix (T-bar) at the AZ center to drive release from several release sites by a common calcium channel cluster. This evolutionary pressure may have been stronger than optimizing postsynaptic amplitude and timing during univesicular release, limiting the number of release sites than can be located at the AZ center.

### 2.5 Unc13A-GluR nano-architecture optimizes synaptic transmission during multivesicular release

Finally, we replicated our previous analyses in the case of multivesicular release. An average synapse was again modeled by assuming 214 postsynaptic GluRs located at an average distance of 121nm from the AZ center. Instead of a single release site, we modeled two release sites that are equally spaced and located on a ring around the AZ center with radius *d* (Fig. 5A). We computed different metrics as a function of *d* (mean mEPSC amplitude, and mean activation time across GluRs *< t*_max_ *>*) and compared the results to the observed distribution of Unc13A positions.

**Figure 5.**
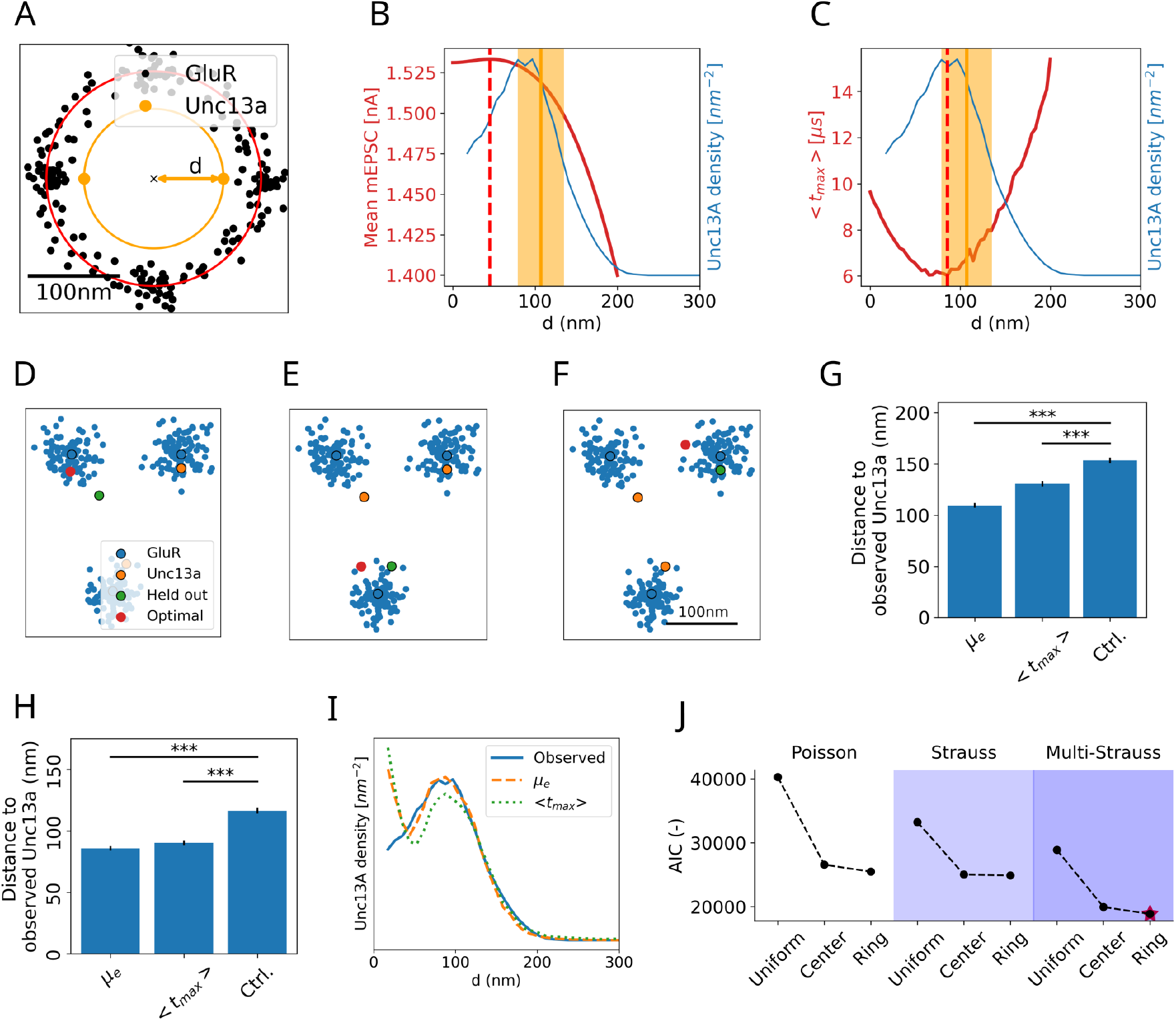
Unc13A-GluR nano-architecture optimizes synaptic transmission during multivesicular release. A: Setting illustration. GluRs were fixed, while 2 Unc13A were located on a ring of variable radius *d*. B-C: Same as Fig. 4H but for multivesicular release (i.e. 2 release sites located on a ring at various distances *d* from the AZ center) instead of a single release site. D-F: Example synapse. The positions of GluRs (blue) were held fixed to their observed values. In this example, there are 3 release sites (orange): at each iteration, one of them was successively held out (green) and the surrogate position that allows to maximize *h* was computed from Eq. 7 (red). See other examples in Supp. Fig. 2. G: The distance between actual and optimal Unc13A (red and green in D-F) was computed for all AZ and compared to a control chance level (i.e. the distances between the held out Unc13A and a random point on the surface of the AZ). H: Same procedure, but optimizing for all *M*_*k*_ Unc13A positions (Eq. 8). I: Comparison of the observed (blue) and optimal (orange and green, Eq. 8) densities of Unc13As as a function of the distance to the AZ center. J: Same as Fig. 1C using optimal (Eq. 9) instead of actual protein positions.

In Fig. 5B-C, the observed density of Unc13A localizations as a function of the distance to the AZ center (blue) is compared to the mean mEPSC amplitude *μ*_*e*_ (Fig. 5B) and activation time *< t*_max_ *>* (Fig. 5C) when release sites are located at different distances *d* from the AZ center. Interestingly, unlike the univesicular case, in the multivesicular setting, the highest *μ*_*e*_ is obtained for *d ≈* 50*nm*, which is closer to the empirically observed distance between Unc13As and AZ center. The remaining offset could be explained e.g. by the presence of VGCC at the AZ center or by the size of Unc13As, which would prevent locating as many proteins in the AZ center as a normative theory would predict. Nevertheless, most observed Unc13A clusters reside within a distance to the AZ center for which *μ*_*e*_ is close to its maximal value, and both empirical and theoretical curves display the same cut-off at *d ≈* 100*nm*. Furthermore, the distance *d* for which the observed Unc13A density is maximal coincides with the theoretical release site distance that minimizes GluR activation time (Fig. 5C). These results suggest that Unc13A-GluR nano-organization optimizes postsynaptic amplitude and activation time during multivesicular synaptic transmission.

We then performed the same numerical optimization procedure for individual synapses as described in Fig. 4E-F. First, we optimized *h* for the position of only one release site at a time: for a given synapse, the positions of all presynaptic Unc13A and postsynaptic GluR clusters are fixed to their observed values, except for one release site (the held-out one). The optimal position for this release site was computed, and compared to the observed held-out Unc13A position. The following computation was performed for the *l*-th release site of the *k*-th active zone:

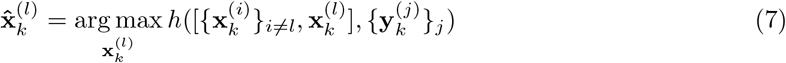

The theoretically optimal position 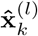 was then compared to the observed Unc13A position 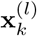. As seen in Fig. 5D-F for a representative synapse with *M*_*k*_ = 3 release sites (blue), the release site position that optimizes *h* (red) closely matches the observed position of the held-out Unc13A cluster (green). This process was repeated across all release sites for all synapses: the mean distance between the optimal and actual Unc13A positions is significantly smaller than chance (i.e., the distance between the observed Unc13A cluster and a random position on the AZ surface) (Fig. 5G: one-sided independent t-test, *p <* 0.001). Using different cost functions for *h* (i.e., the mean mEPSC amplitude *μ*_*e*_, or the mean activation time *< t*_max_ *>*) yields similar results.

Next, we performed the same analysis, but optimizing the positions of all release sites at once. If the number of release sites at the *k*-th AZ is *M*_*k*_, then the *M*_*k*_ positions were optimized (the GluR positions 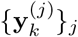 being fixed to their observed positions, as before):

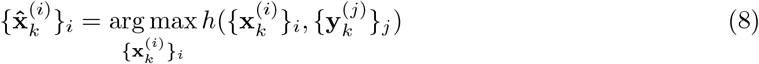

The distance between the observed Unc13A cluster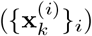 and the optimal release site 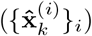 is again significantly smaller than chance (Fig. 5H: one-sided independent t-test, *p <* 0.001), suggesting that the Unc13A cluster locations maximize the activation of postsynaptic GluRs during multivesicular release. Furthermore, there is a good qualitative match between the observed and theoretically predicted spatial Unc13A distributions (Fig. 5I).

Finally, we closed the loop by performing the same model-selection procedure as in Section 2.1, but applying it to the theoretically optimal positions for Unc13A and GluRs 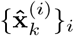 and 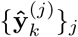 instead of the observed ones 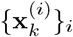 and 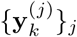. The former were computed by numerically optimizing *h*, under the constraints of the same minimum distance between the observed clusters (Fig. 4I):

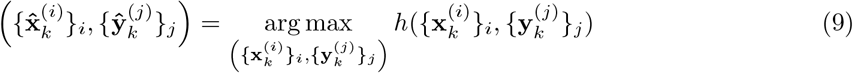

subject to

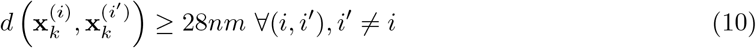

and

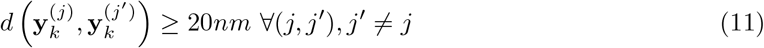

We thus simultaneously optimized the positions of all pre- and postsynaptic proteins, the only constraints being the observed minimal Unc13A-to-Unc13A and GluR-to-GluR distances (respectively 28*nm* and 20*nm*), to avoid overlapping proteins. In particular, no constraints on density intensity nor proteins interaction (such as a type-dependent interaction radius) were set, in order not to bias model selection. Proteins were initialized at their observed positions to represent prior knowledge on structural constraints. Both cases yielded the same conclusion (compare Fig. 5J to Fig. 1C): the model with the best goodness-of-fit is the one with ring-like intensity function patterns, negative interaction between protein cluster locations of the same type and positive interactions between proteins of different types. This further highlights that the observed distribution of synaptic proteins is close to the one that optimizes the amplitude and timing of synaptic transmission at the NMJ during multivesicular release.

## 3 Discussion

We here investigated the organization of synaptic nano-architecture and its relationship to synaptic transmission at the *Drosophila* NMJ using a combination of model-based approaches, super-resolution microscopy, and electrophysiology. Model-based quantification supports non-randon, transsynaptically-aligned rings of clusters formed by the presynaptic protein Unc13A and post- synaptic GluR receptors. Our analytical model predicts that most receptors are activated by the neurotransmitter released from one synaptic vesicle during univesicular release (UVR), and that release from the AZ center optimizes for postsynaptic current amplitude, in contradiction to observed Unc13A locations. However, release from the observed Unc13A locations is predicted to optimize for postsynaptic amplitude during multivesicular release (MVR).

Our findings challenge the prevailing model that transsynaptic nano-architecture primarily optimizes the amplitude of spontaneous mEPSCs or AP-evoked synaptic transmission [8, 16]. Specifically, our model-based approach does not support amplitude optimization during UVR at the Drosophila NMJ, given that observed Unc13A locations are offset from the AZ center positions predicted to be optimal for postsynaptic activation. Rather, our model predicts the activation of the majority of GluRs within the PSD during UVR, consistent with electrophysiology-based estimates of 200 activated receptors per release event [32]. By contrast, the transsynaptically-aligned ring organization of Unc13A and GluRs is predicted to optimize postsynaptic amplitude specifically during MVR. Given that MVR is predominantly observed during conditions of enhanced presynaptic release at the *Drosophila* NMJ, our data suggest that transsynaptic nano-architecture may be adapted to elevated presynaptic release states and synaptic plasticity rather than to baseline UVR. Several lines of reasoning may account for the offset between optimal and observed Unc13A positions during UVR. First, structural constraints may represent an evolutionary compromise: the concentration of voltage-gated calcium channels together with scaffolding proteins such as Brp and RIM-binding protein at the AZ center enables a common calcium channel cluster to drive release from multiple sites, an architectural logic that may have exerted stronger evolutionary pressure than optimizing postsynaptic amplitude during individual release events. Second, the spatial distribution of Unc13A is not uniform; a significant fraction localizes to the AZ center, which may sustain release during UVR, while the peripheral pool, peaking at approximately 75 nm from the AZ center, may selectively support MVR. Third, it cannot be excluded that a subset of Unc13A clusters does not correspond to functional release sites, contributing to the apparent positional offset. Fourth, and more broadly, synaptic nano-architecture may optimize not for amplitude and timing per se, but for synaptic plasticity, for the coordinated targeting of multiple release sites by a shared calcium channel cluster, or for the molecular signaling capacity required for the modulation of synaptic transmission. Consistent with a role in plasticity, amplitude optimization during MVR may be particularly relevant during presynaptic homeostatic potentiation, a form of synaptic plasticity during which quantal size is reduced and MVR is enhanced to compensate for diminished postsynaptic sensitivity.

Future studies will be required to directly test these models, in particular to resolve the functional contributions of distinct Unc13A subpopulations, to determine the conditions under which MVR predominates at individual AZs, and to examine whether the nano-architectural principles identified here are conserved across synaptic types and species.

## 4 Material and methods

### 4.1 Immunohistochemistry

*Drosophila* stocks were stored in an incubator at 25°C, in plastic tubes with food containing molasses and closed with cotton wool. Experiments were made on 3rd instar larvae freely moving on the tube wall. The *w*^118^ strain was used for all the experiments.

**Day 1**

#### Dissection

Moving 3rd instar larvae without horns were selected and placed on a solid agar-gel staining petri-dish plate. Larva was placed so that the back side and the trachea were facing up and therefore visible. The larva head and tail were pin down so that it was fixed to the plate and three drops of HL3 were added on top of the larva. The larva was then cut longitudinally and the organs were removed. Next, two pins were placed on each side at the upper and lower extremities, so that the larva was fixed as an open book. HL3 was washed out and replaced with new HL3.

#### Fixation

Once in open book preparation, HL3 was removed and the larva was fixed with Ethanol for 15 minutes on ice. Ethanol was then removed, and the larva quickly washed three times with PBS. Pins were carefully removed, and the larva placed in a 1mL Eppendorf containing 1mL of PBST. The Eppendorf was placed on a rotating wheel at room temperature for 50 minutes, while PBST was replaced by new PBST every 10 minutes (5 times during the washing period). After the 50 minutes, PBST was replaced by blocking solution containing 3% NGS and the Eppendorf left on the rotating wheel for 2 hours at room temperature. At the end of the 2 hours, blocking solution was removed and replaced by the primary antibody solution, and the Eppendorf was left incubating in a rotating wheel at 4°C overnight.

**Day 2**

On the next day, the primary solution was removed and the larva washed 5x 10 min in PBST as described above. Starting at the beginning of the washing and until mounting the slide, the Eppendorf was covered with aluminum foil to protect from the light. After the washing period, PBST was removed and the secondary antibody solution was added, and the tube left rotating at room temperature for 2 hours. The secondary antibody solution was removed and the larva washed 5x 10minutes in PBST as previously. Then, the larvae were mounted on a glass slide (Thermo Scientific, Menzel Gläser. 76×26mm) with Prolong gold mounting medium and coveslipped (Menzel Gläser, 24×50mm, #1.5).

Solution preparations:

- PBS (1x): Bottle made from the kitchen of the Department of Molecular Life Science, University of Zurich.
- PBST: 0.1% TritonX-100 in PBS
- Blocking solution: 3% NGS in PBST (0.1% Triton)

Primary antibodies:

- anti-Brp (nc82), mouse, 1:100;
- anti-GluRIIC, rabbit, 1:100.

Secondary antibodies:

- Abberior STAR 635 P, anti-rabbit (Sigma-Aldrich), 1:100;
- Atto 594, anti-mouse (Sigma-Aldrich), 1:100.

### 4.2 STED microscopy

#### Image acquisition

Images were acquired with a time-gated stimulated emission depletion (STED) microscope based on a Leica SP8 inverted confocal microscope (Leica SP8 3X) of UZH’s Center for Microscopy and Image Analysis (ZMB). Images were taken using a HC PL APO 100x/1.40 Oil STED WHITE objective with immersion oil conforming to ISO 8036 with a diffraction index of *n* = 1.5180. The confocal channels were used to position the sample and to set up the Z-stack range. The Z-step size was 0.13*μm*. Channel 0 and 2 were used to image the STAR 635P antibody whereas channel 1 and 3 the ATTO 594 antibody. For each staining before imaging, the saturation level for each channel was checked and the laser power adjusted accordingly. In addition, a pixel size of 20nm was used.

The following settings for each channel were used:

- Channel 0: white at 640 nm with 27.6% power, STED 775nm laser with 0% power, HyD in counting mode with detection from 646 to 714, no gating, line accumulation 1.
- Channel 1: white at 594 nm with 15.5% power, STED 775nm laser with 0% power, HyD in counting mode with detection from 599 to 632, no gating, line accumulation 1.
- Channel 2: white at 640 nm with 27.5% power, STED 775nm laser with 76.5% power, HyD in counting mode with detection from 646 to 714, gating from 0.7 to 6ns, line accumulation 6.
- Channel 3: white at 594 nm with 31.4% power, STED 775nm laser with 63.5% power, HyD in counting mode with detection from 599 to 632, gating from 0.7 to 6ns, line accumulation 6.

Images were acquired with LAS X (Leica Application Suite X, version 2.0; Leica Microsystems, Germany).

#### Deconvolution

For image deconvolution, Huygens Professional was used (Huygens compute engine 20.04, Scientific Volume Imaging B.V., The Netherlands). We used the automatically generated point spread function (PSF), and the Auto Stabilize features for lateral drift correction. Furthermore, we used the automatic detection, with lowest estimation mode and an area radius of 0.7 micron for the background detection, as well as the Good’s Roughness Maximum Likelihood Estimation (GMLE) deconvolution algorithm with the following default estimated parameters: maximum iterations of 10, signal to noise ratio of 15 for confocal channels and 7 for STED channels (might vary as the SNR estimator could give different values depending on the image) and a quality threshold of 0.003 for confocal channels and 0.005 for STED channels.

### 4.3 GluR quantification via dSTORM

For dSTORM imaging of GluRs, HA-tagged GluRIIA was expressed from a genomic transgene in the GluRIIA and GluRIIB double mutant background (dfclh4/A22; GluRIIA^*HA*^/+) as described previously [38].

Third-instar larvae were prepared, stained, and imaged according to established methods [36]. In brief, we used the primary antibodies rabbit-*α*-HA (Cell Signaling, AB 1549585) and mouse- *α*-Brp [nc82, AB 2314866]. The secondary antibodies were goat-*α*-mouse AlexaFluor 532 (Life Technologies, A11002, AB 2534070), goat-*α*-rabbit AlexaFluor532 (Life technologies, AB 2534076), and goat-*α*-rabbit F(ab’)2 fragments (Invitrogen, AB 429691) labelled with Cy5. The samples were embedded in photoswitching buffer consisting of 100 mM *β*-mercaptoethylamine, oxygen scavenger system (1 % [wt/vol] glucose, 5 U/ml glucose oxidase and 100 U/ml catalase [39]), pH 8.0, and imaged on an inverted microscope (Axio Observer Z1), equipped with a focus corrector (Definite Focus, Zeiss), an oil-immersion objective (alpha plan-apochromat, 100x, 1.46 N.A., Zeiss), and EMCCD cameras (iXON Ultra 897, Andor Technology). Excitation was performed with 532 nm (Laser Quantum gem) and 640nm (iBeam smart, TOPTICA Photonics) lasers. During acquisition, 15,000 frames were captured with 10 ms integration time. The super-resolution images were reconstructed using the ThunderSTORM Fiji plugin, a subpixel binning of 10nm/pixel was applied, and two-color images were aligned using the bUnwarpJ Fiji plugin. 2D histograms were reconstructed using the localization tables generated by ThunderSTORM and then analyzed using custom Fiji macros to quantify GluRs.

The number of GluRs per postsynaptic density (PSD) was estimated in principle as previously described [36]. In order to measure *L*_*E*_, the number of localizations presented per primary antibody (ab1) bound to Cy5-labelled secondary antibody (ab2) within the relevant subcellular nano-environment, the preparations were stained with a very low concentration of ab1 (1:80,000; ab2: 5.2 x 10^*−*8^ M Cy5, 100 %) together with an antibody against Brp (mouse-*α*-Brp/nc82, 1:100, labelled by goat-*α*-mouse AlexaFluor 532, 1:500). The Brp label served as a proxy for the adjacent PSD and only individual spots within the region defined by Brp were analyzed.

The saturation of ab1 was assessed via stepwise dilutions of ab1 (1:20 to 1:40,000) while maintaining a constant concentration of ab2 (5.2 x 10^*−*8^ M) with a fixed ratio of Cy5 and AF532 (10 % Cy5, 90 % AF532). For these experiments, the concentration of Cy5 F(ab’)2 fragments was reduced to ensure the separation of individual fluorescence emissions at high ab1 concentrations. The average number of localizations per PSD was plotted against the ab1 dilution and fitted with a 4-parameter logistic function to obtain an estimate of the maximum localization value, *L*_1_.

Using the localization values in Table 2 to solve the equation

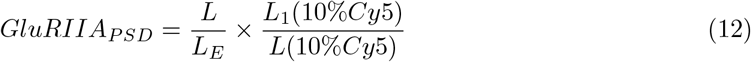

gives an estimate of 214.6 *±* 65 GluRs s.e.m. per PSD.

**Table 2:** Localization values from antibody titrations. *L*: localizations at experimental conditions (100 % Cy5, 1:2,000 ab1). *L*_*E*_: localizations corresponding to single primary antibody (100% Cy5, 1:80,000 ab1). *L*_1_ (10% Cy5): localizations at saturation of ab1 (1:20), low Cy5 concentration. *L* (10% Cy5): localizations at experimental conditions of ab1 (1:2,000), low Cy5 concentration.

| | $L$ | $L_E$ | $L_1$ (10 % Cy5) | $L$ (10 % Cy5) |
| --- | --- | --- | --- | --- |
| Localizations | 2862.4 | 31.3 | 1583.4 | 675.7 |
| s.e.m | 68 | 1.4 | 216.5 | 178.6 |

### 4.4 Gibbs Point Processes

The probability distribution for the positions of a set of observations **X, Y** is computed as

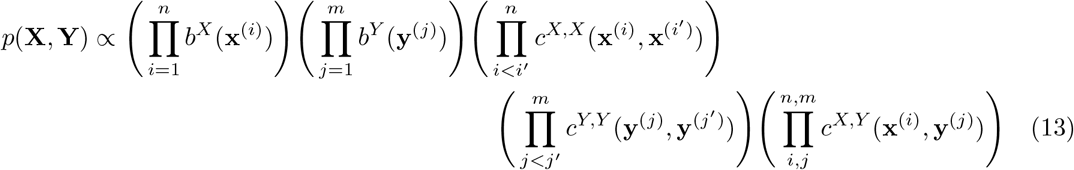

where 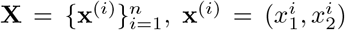 are the 2D coordinates of *n* Unc13A clusters and 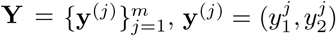are the 2D coordinates of *m* GluR clusters.

First, point processes are characterized by their intensity function *b*^*z*^(**u**) (i.e. the density of points), which can be either uniform or be a function of the position **u** = (*x*_1_, *x*_2_) in the 2D plane and of the type *z* of points, with *z ∈ {X, Y }*. We propose 3 different intensity functions. First, a uniform one:

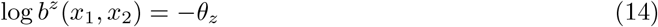

Under the uniform hypothesis, the process is said to be homogeneous (i.e. the point density is the same over the whole surface of the AZ plane). Second, a distance-to-center intensity function:

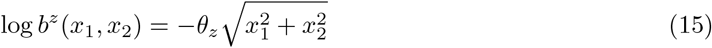

Under this hypothesis, the density of proteins is assumed to be a decreasing function of the distance to the center of the AZ plane. Finally, a ring patterns intensity function:

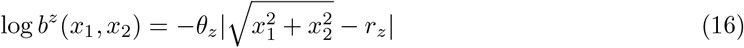

where *r*_*z*_ is the mean distance of proteins of type *z* to the center of the AZ plane. Indeed, Unc13A and GluR have been observed to be organized as ring-like patterns, with an inner ring of Unc13A proteins close to the center of the AZ and an outer ring of GluR clusters (Figure 1A). Performing model selection would precisely allow to verify this hypothesis, by confronting its goodness-of-fit to that of other models.

Second, point processes are characterized by the interpoint interactions, i.e. by possible dependencies between the points’ positions and types. The functions 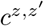correspond to pairwise interaction terms, which are symmetric between positions **u** and **v** (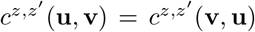.

Here, we propose 3 different interaction schemes. First, a Poisson point process:

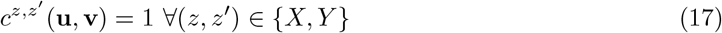

In a Poisson point process, all points are i.i.d., irrespective of their types. Second, a Strauss point process:

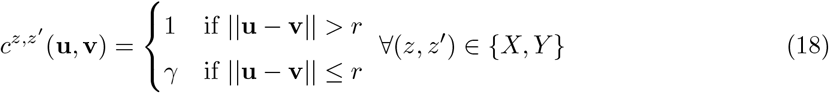

A Strauss process specifies an interaction radius *r* between two points **u** and **v**. It reduces to a Poisson process for *γ* = 1 and to a Hardcore process for *γ* = 0 (i.e. a hard exclusion radius around each protein). Finally, a multi-Strauss point process:

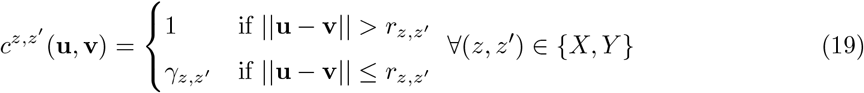

It further generalizes the Strauss process (Eq. 18) to be type-dependent: the probability parameter *γ* and the interaction radius *r* will depend on the respective types of **u** and **v** (*Unc13A-to-Unc13A, Unc13A-to-GluR*, or *GluR-to-GluR*).

Procedures for likelihood computation for GPP have been previously described in [40] and [41], while maximum likelihood estimators are computed using the Berman-Turner algorithm [42].

### 4.5 Model comparison

The AIC is a model selection score that compares candidate models based on their ability to explain observations while penalizing their number of free parameters [22–24]. For a given model *M* and a set of observations *D*, it is defined as:

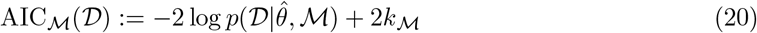

where 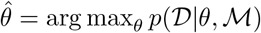 is the maximum likelihood estimator of the parameters of *M* and *k*_*M*_ is its number of free parameters. The first term represents the ability of the model *M* to explain *D*, while the penalty term *k*_*M*_ favors simpler models and prevents overfitting: the lower the AIC, the better the model [43].

### 4.6 Statistical model of glutamate spread and postsynaptic receptor activation

Assuming that a spontaneous release happened at one of the presynaptic release sites, we use the following notations: Δ*S* is the surface of a postsynaptic receptor (expressed in *nm*^2^); *A* is the total surface of the active zone; *n* is the number of released glutamate molecules; and *ρ* is their destiny per surface unit, 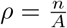.

The probability *p* that a single receptor will be activated can be approximated as the probability that at least one of the *n* molecules will be in the same area as the receptor (assuming a perfect affinity between glutamate and receptors):

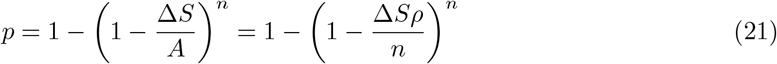

Assuming that *n* is large yields

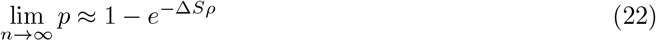

Substituting Eq. 3 into 22 finally yields

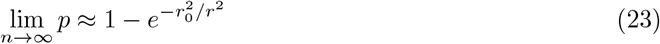

with

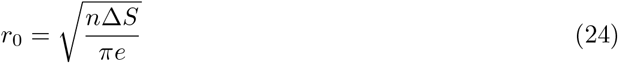

### 4.7 Extension to multivesicular release

Glutamate concentrations from different release sites are additive: for *M* release sites, the maximum glutamate concentration that will be reached at a given point is

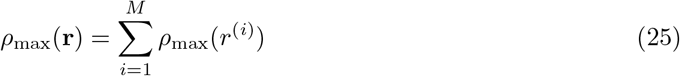

where 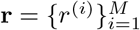 are the distances between this point and the *M* release sites.

Eq. 4 readily extends to multi-vesicular release, i.e. to compute the activation probability of a postsynaptic receptor located at distances 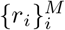from *M* release sites. Indeed, it can be expressed as the composition of 2 functions:

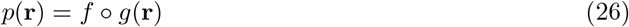

where 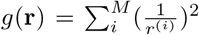 computes the glutamate concentration as a function of the distances between proteins (since concentration adds up, in line with Eq. 25) and 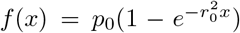 transforms this observed concentration into an activation probability (in line with Eq. 4).

### 4.8 Generative model of postsynaptic currents

We propose a generative model for mEPSCs with the following notations: mEPSCs were recorded at a given NMJ, which corresponds to a single axon with several active zones, all linked to the same muscle cell. We thus denote 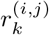the distance between release site 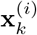 and postsynaptic receptor 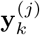 at the *k*-th Active Zone of the studied NMJ:

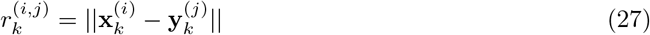

Empirical samples for 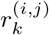 can be obtained from the observed positions of presynaptic Unc13A and postsynaptic GluRs (Section 2.1). 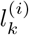 is the number of receptors activated by a release at site 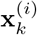:

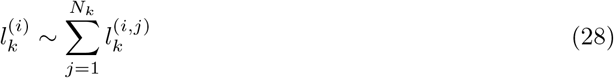

where 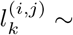 Bernoulli 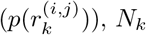is the number of GluR at the *k*-th synapse, and *p*(·) is the sensitivity function defined by Eq. 4. *N*_*k*_ is assumed to be different for each synapse and to follow a normal distribution with mean *μ*_*N*_ and variance 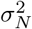

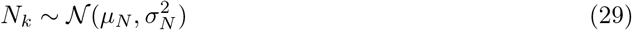

We further assume that the recorded postsynaptic current is the sum of the currents elicited by each activated receptor [3, 25]. The postsynaptic current elicited by the activation of one receptor is modeled to follow a Gamma distribution with shape and scale parameters *α* and *β*, or equivalently with a mean *q*_0_ (representing the mean postsynaptic current elicited by the activation of one GluR) and a standard deviation *σ* (representing effects that are not accounted for in the model, such as recording noise or different dynamics between GluRIIA and GLuRIIB subunits) with

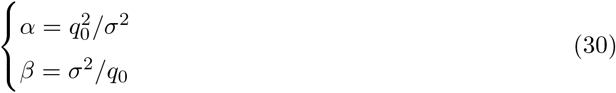

The activation of 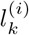independent receptors will thus elicit a postsynaptic current 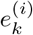drawn from

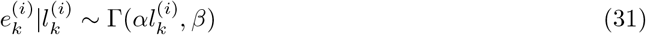

Finally, we respectively denote as *I* = *i* and *K* = *k* the events corresponding to the postsynaptic current being elicited by a glutamate release at the *i*-th release site of the *k*-th AZ. The final probability density function for a postsynaptic current event *e* is finally computed by marginalizing the joint probability *p*_*θ*_(*e, K, I, l*):

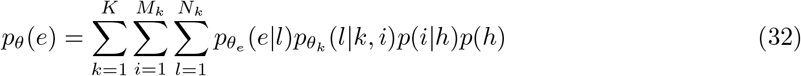

where *p*_*θ*_(*e*|*l*) is given by Eq. 31, *p*_*θ*_(*l*|*k, i*) is given by Eq. 28, *p*(*i*|*k*) = 1*/M*_*k*_, and *p*(*k*) = 1*/K* (i.e. each release site from all AZ are considered equally likely to have generated the observed mEPSC). *θ* corresponds to the vector of parameters: *θ* = [*θ*_*e*_, *θ*_*l*_] with *θ*_*e*_ = [*q*_0_, *σ*] (Eq. 30) and *θ*_*l*_ = [*μ*_*N*_, *σ*_*N*_, *r*_0_, *p*_0_] (Eqs. 4 and 29).

Note that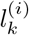 follows a Poisson-Binomial distribution (Eq. 28), which probability mass function becomes intractable unless the number of trials is small [44, 45]. The marginal sum over *l* in Eq. 32 is thus represented by the following point-based estimation:

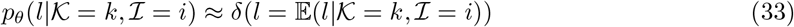

with

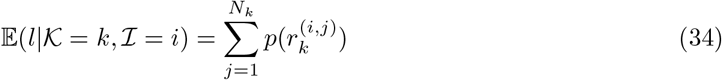

Note that the distribution of mEPSCs amplitudes (Fig. 3A and Eq. 32) is computed from the observed positions of Unc13A and GluRs (Eq. 27): currents are assumed to be elicited by a single fusion event taking place at one of the observed release sites, i.e. at one of the observed Unc13A positions (Fig. 1A). Each active zone has different Unc13A and GluR positions, hence eliciting different postsynaptic amplitudes for different positions of the release event. This variability across active zones and release site positions underlies the variability of postsynaptic current amplitudes.

Following Eq. 34, the mean mEPSC *μ*_*e*_ is computed as

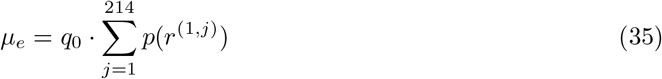

In Eq. 35, the term *r*^(1,*j*)^ refers to the distance between the position of the considered release site and the *j*-th GluR.

## Acknowledgments

We thank Denis-Gabriel Caprace, Jakob Jordan, and Igor Delvendahl for the fruitful discussions. The funders played no role in study design, data collection, analysis and interpretation of data, or the writing of this manuscript.

## 5 Supplementary material

**Supplementary Figure 1:**
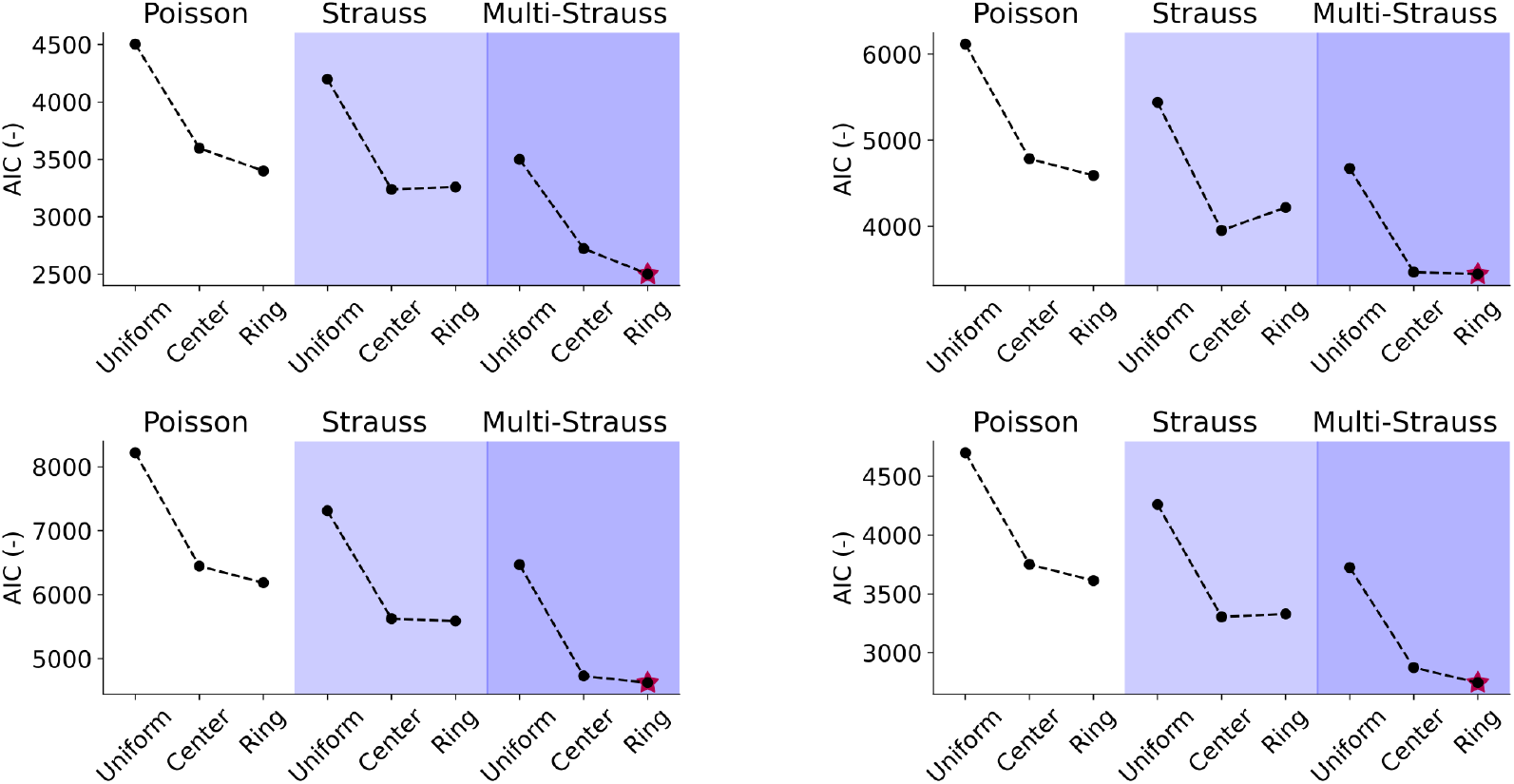
Same as Fig. 1C for the other 4 recorded NMJs.

**Supplementary Figure 2.**
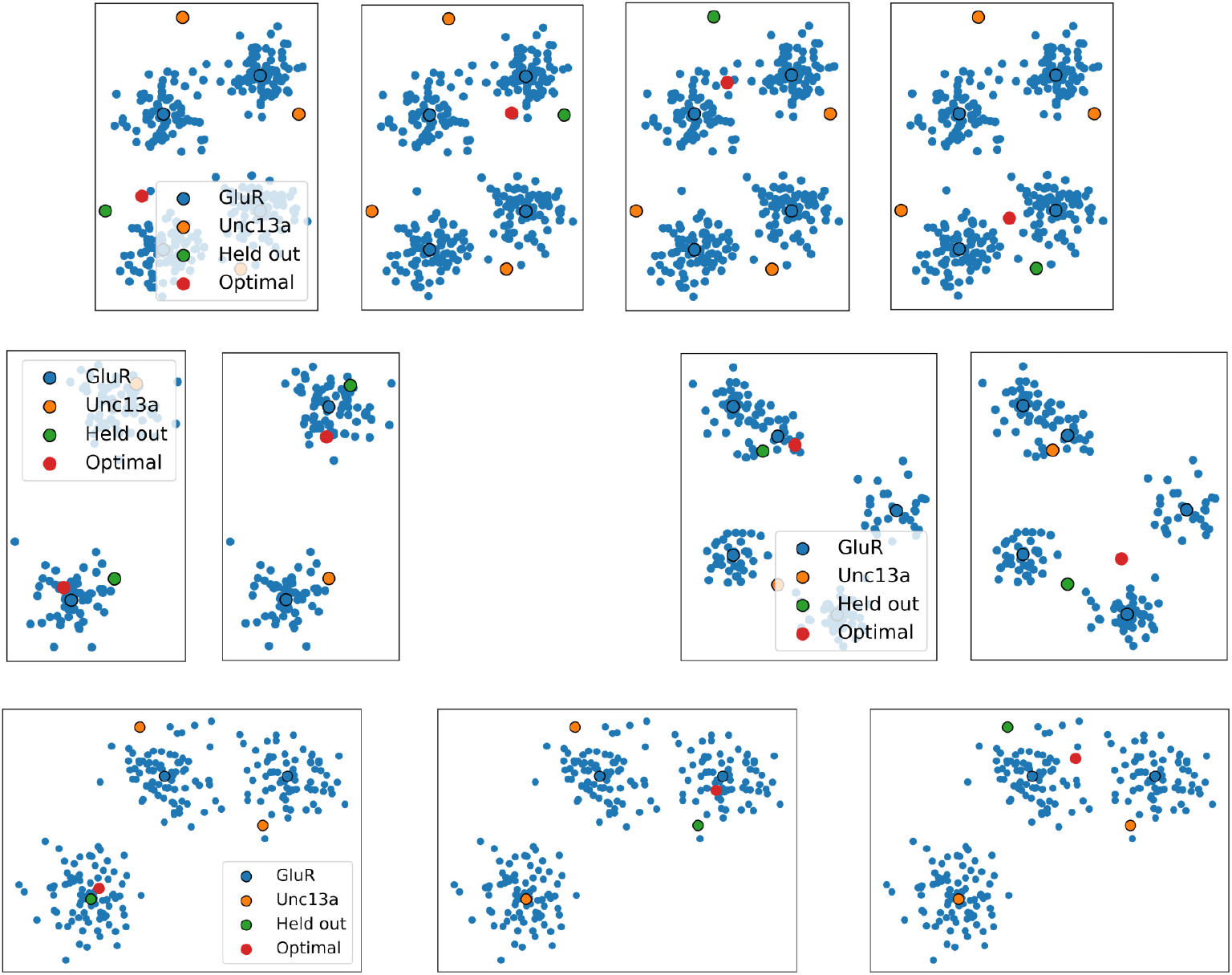
Same as Fig. 5D-F for 4 other examples synapses, with various numbers of Unc13A and GluRs.

## References

[1] M. Heckmann, F. Parzefall, and J. Dudel, “Activation kinetics of glutamate receptor channels from wild-type drosophila muscle,” Pflügers Archiv-European Journal of Physiology, vol. 432, no. 6, pp. 1023–1029, 1996.

[2] M. Heckmann and J. Dudel, “Recordings of glutamate-gated ion channels in outside-out patches from drosophila larval muscle,” Neuroscience letters, vol. 196, no. 1-2, pp. 53–56, 1995.

[3] S. Raghavachari and J. E. Lisman, “Properties of quantal transmission at ca1 synapses,” Journal of neurophysiology, vol. 92, no. 4, pp. 2456–2467, 2004.

[4] V. V. Uteshev and P. S. Pennefather, “A mathematical description of miniature postsynaptic current generation at central nervous system synapses,” Biophysical Journal, vol. 71, no. 3, pp. 1256–1266, 1996.

[5] X. Li, G. Hémond, A. G. Godin, and N. Doyon, “Computational modeling of trans-synaptic nanocolumns, a modulator of synaptic transmission,” Frontiers in Computational Neuroscience, vol. 16, p. 969 119, 2022.

[6] D. Atasoy et al., “Spontaneous and evoked glutamate release activates two populations of nmda receptors with limited overlap,” Journal of Neuroscience, vol. 28, no. 40, pp. 10 151–10166, 2008.

[7] K. M. Franks, C. F. Stevens, and T. J. Sejnowski, “Independent sources of quantal variability at single glutamatergic synapses,” Journal of Neuroscience, vol. 23, no. 8, pp. 3186–3195, 2003.

[8] A. M. Ramsey et al., “Subsynaptic positioning of ampars by lrrtm2 controls synaptic strength,” Science Advances, vol. 7, no. 34, eabf3126, 2021.

[9] H. Sakamoto et al., “Synaptic weight set by munc13-1 supramolecular assemblies,” Nature neuroscience, vol. 21, no. 1, pp. 41–49, 2018.

[10] K. S. Liu et al., “Rim-binding protein, a central part of the active zone, is essential for neurotransmitter release,” Science, vol. 334, no. 6062, pp. 1565–1569, 2011.

[11] R. Kittel, S. Hallermann, S. Thomsen, C. Wichmann, S. Sigrist, and M. Heckmann, Active zone assembly and synaptic release, 2006.

[12] R. J. Kittel et al., “Bruchpilot promotes active zone assembly, ca2+ channel clustering, and vesicle release,” Science, vol. 312, no. 5776, pp. 1051–1054, 2006.

[13] A.-H. Tang, H. Chen, T. P. Li, S. R. Metzbower, H. D. MacGillavry, and T. A. Blanpied, “A trans-synaptic nanocolumn aligns neurotransmitter release to receptors,” Nature, vol. 536, no. 7615, pp. 210–214, 2016.

[14] P. Muttathukunnel, P. Frei, S. Perry, D. Dickman, and M. Müller, “Rapid homeostatic modulation of transsynaptic nanocolumn rings,” Proceedings of the National Academy of Sciences, vol. 119, no. 45, e2119044119, 2022.

[15] M. A. Böhme et al., “Active zone scaffolds differentially accumulate unc13 isoforms to tune ca2+ channel–vesicle coupling,” Nature neuroscience, vol. 19, no. 10, pp. 1311–1320, 2016.

[16] K. T. Haas et al., “Pre-post synaptic alignment through neuroligin-1 tunes synaptic transmission efficiency,” elife, vol. 7, e31755, 2018.

[17] A. Scimemi and M. Beato, “Determining the neurotransmitter concentration profile at active synapses,” Molecular neurobiology, vol. 40, no. 3, pp. 289–306, 2009.

[18] S. Li et al., “Asynchronous release sites align with nmda receptors in mouse hippocampal synapses,” Nature communications, vol. 12, no. 1, pp. 1–13, 2021.

[19] A. A. Faisal, J. A. White, and S. B. Laughlin, “Ion-channel noise places limits on the miniaturization of the brain’s wiring,” Current Biology, vol. 15, no. 12, pp. 1143–1149, 2005.

[20] W. M lynarski, M. Hledík, T. R. Sokolowski, and G. Tkačik, “Statistical analysis and optimality of neural systems,” Neuron, vol. 109, no. 7, pp. 1227–1241, 2021.

[21] M. Galkov, P. Nemcova, D. Dietrich, and S. Schoch, “Active zone: Linking resolution levels of microscopic modalities,” Nano-organization of the Synapse: From Structure to Function, pp. 79–106, 2026.

[22] H. Akaike, “A new look at the statistical model identification,” IEEE transactions on automatic control, vol. 19, no. 6, pp. 716–723, 1974.

[23] K. P. Burnham and D. R. Anderson, “Multimodel inference: Understanding aic and bic in model selection,” Sociological methods & research, vol. 33, no. 2, pp. 261–304, 2004.

[24] D. Anderson and K. Burnham, “Model selection and multi-model inference,” Second. NY: Springer-Verlag, vol. 63, no. 2020, p. 10, 2004.

[25] E. Tarusawa et al., “Input-specific intrasynaptic arrangements of ionotropic glutamate receptors and their impact on postsynaptic responses,” Journal of Neuroscience, vol. 29, no. 41, pp. 12 896–12 908, 2009.

[26] D. Nair et al., “Super-resolution imaging reveals that ampa receptors inside synapses are dynamically organized in nanodomains regulated by psd95,” Journal of Neuroscience, vol. 33, no. 32, pp. 13 204–13 224, 2013.

[27] S. Masoli, M. F. Rizza, M. Tognolina, F. Prestori, and E. D’Angelo, “Computational models of neurotransmission at cerebellar synapses unveil the impact on network computation,” Frontiers in Computational Neuroscience, 2022.

[28] H. Zhan, J. Bruckner, Z. Zhang, and K. O’Connor-Giles, “Three-dimensional imaging of drosophila motor synapses reveals ultrastructural organizational patterns,” Journal of neurogenetics, vol. 30, no. 3-4, pp. 237–246, 2016.

[29] F. Ventriglia, “Effect of filaments within the synaptic cleft on the response of excitatory synapses simulated by computer experiments,” Biosystems, vol. 104, no. 1, pp. 14–22, 2011.

[30] X. Li, G. Hémond, A. G. Godin, and N. Doyon, “Computational modeling of trans-synaptic nanocolumns, a modulator of synaptic transmission,” Frontiers in Computational Neuroscience, vol. 16, 2022, issn: 1662-5188. doi: 10.3389/fncom.2022.969119. [Online]. Available: https://www.frontiersin.org/articles/10.3389/fncom.2022.969119.

[31] C. E. Jahr, “High probability opening of nmda receptor channels by l-glutamate,” Science, vol. 255, no. 5043, pp. 470–472, 1992.

[32] A. DiAntonio, S. A. Petersen, M. Heckmann, and C. S. Goodman, “Glutamate receptor expression regulates quantal size and quantal content at the drosophila neuromuscular junction,” Journal of Neuroscience, vol. 19, no. 8, pp. 3023–3032, 1999.

[33] M. Baccino-Calace, K. Schmidt, and M. Müller, “The e3 ligase thin controls homeostatic plasticity through neurotransmitter release repression,” Elife, vol. 11, e71437, 2022.

[34] P. S. O’Neill et al., “A deep learning framework for automated and generalized synaptic event analysis,” ELife, vol. 13, RP98485, 2025.

[35] A. Weyhersmüller, S. Hallermann, N. Wagner, and J. Eilers, “Rapid active zone remodeling during synaptic plasticity,” Journal of Neuroscience, vol. 31, no. 16, pp. 6041–6052, 2011.

[36] N. Ehmann et al., “Quantitative super-resolution imaging of bruchpilot distinguishes active zone states,” Nature communications, vol. 5, no. 1, p. 4650, 2014.

[37] C. Feeney, S. Karunanithi, J. Pearce, C. Govind, and H. Atwood, “Motor nerve terminals on abdominal muscles in larval flesh flies, sarcophaga bullata: Comparisons with drosophila,” Journal of Comparative Neurology, vol. 402, no. 2, pp. 197–209, 1998.

[38] T. M. Rasse et al., “Glutamate receptor dynamics organizing synapse formation in vivo,” Nature neuroscience, vol. 8, no. 7, pp. 898–905, 2005.

[39] P. Schäfer S. Van De Linde, J. Lehmann, M. Sauer, and S. Doose, “Methylene blue-and thiol-based oxygen depletion for super-resolution imaging,” Analytical chemistry, vol. 85, no. 6, pp. 3393–3400, 2013.

[40] A. Baddeley et al., “Analysing spatial point patterns in r,” in Workshop notes version, vol. 3, 2008.

[41] A. Baddeley, E. Rubak, and R. Turner, Spatial point patterns: methodology and applications with R. CRC press, 2015.

[42] M. Berman and T. R. Turner, “Approximating point process likelihoods with glim,” Journal of the Royal Statistical Society: Series C (Applied Statistics), vol. 41, no. 1, pp. 31–38, 1992.

[43] C. Gontier, “Statistical approaches for synaptic characterization,” Ph.D. dissertation, University of Bern, 2022.

[44] S. X. Chen and J. S. Liu, “Statistical applications of the poisson-binomial and conditional bernoulli distributions,” Statistica Sinica, pp. 875–892, 1997.

[45] M. Fernández and S. Williams, “Closed-form expression for the poisson-binomial probability density function,” IEEE Transactions on Aerospace and Electronic Systems, vol. 46, no. 2, pp. 803–817, 2010.

